# Flagellar toxicity: flagellar synthesis is lytic for *Bacillus subtilis* in the absence of PBP1

**DOI:** 10.64898/2026.05.21.726928

**Authors:** Caroline Dunn, Kehinde O. Adebiyi, Daniel B. Kearns

## Abstract

Flagella are large transenvelope nanomachines but how they transit the peptidoglycan in Gram positive bacteria is poorly understood. A recent model suggested that flagellar basal bodies diffuse in the membrane and become captured at locations in the peptidoglycan with a pore diameter that could accommodate the axle-like flagellar rod. Mutation of penicillin binding protein 1 (PBP1/PonA), a cell wall repair protein thought to decrease peptidoglycan pore frequency and/or size, resulted in a severe growth defect and cell lysis in the ancestral strain of *Bacillus subtilis* that was dependent on flagellar synthesis. Genetic analysis indicated that toxicity was due to completion of the flagellar hook, which activated the flagellar sigma factor SigD. SigD, in turn, activated a suite of peptidoglycan hydrolases that caused cellular lysis when PBP1 was absent. In addition, mutations that resulted in high levels of the stress response factor Spx could lessen the toxicity, while PBPX, a putative teichoic acid D-alanylase, was required for autolysis. In sum our results indicate that flagellar synthesis, not normally associated with cell viability, causes cell wall stress and under some conditions, cell death. Moreover, our work indicates that cost of envelope integrity by flagellar synthesis may be underappreciated due to strain domestication, and suggests that specialized systems may compensate for the cost of assembly of transenvelope machines in general.

**SIGNIFICANCE:** Bacteria assemble nanomachines through the cell envelope but how the machines transit the peptidoglycan is poorly understood. Here we find that assembly of trans-envelope flagella results in cell lysis of *Bacillus subtilis* when the peptidoglycan repair protein PBP1 is absent. Lysis was due to multiple peptidoglycan lyases expressed as a consequence of flagellar assembly, and lytic activity required another PBP homolog, PBPX. Our work indicates that flagella, not normally thought to impact cell viability, can be lethal at the level of cell envelope integrity.

## INTRODUCTION

The bacterial cytoplasm has a high concentration of solutes, and water flows continuously into the cell in hypotonic environments. To protect the plasma membrane from catastrophic osmotic hyper-expansion, bacteria construct an extracellular wall of peptidoglycan that forms a continuous mesh of polysaccharide chains crosslinked by amino acids (1). The average pore diameter of peptidoglycan has been measured to be 2-4 nm, a porosity less than the calculated minimum of 8 nm necessary to contain the plasma membrane under high osmotic pressure (2,3). For cells to survive, the PG superstructure must be unfailingly confluent and stable, while also accommodating dynamic synthesis and remodeling during cell growth and division (4). Moreover, bacteria assemble large protein complexes such as secretion apparati and motility machines that transverse the envelope. In many cases, the diameter of the transenvelope complexes is substantially wider than the calculated average PG pore size, but how the geometric incompatibility is resolved at the molecular level is poorly understood (5–8). Arguably the best studied trans-peptidoglycan machine is the bacterial flagellum.

Bacterial flagella are assembled from the inside-out and span all layers of the cell envelope (9,10). The first component built is the basal body that sits in the plasma membrane and houses a type III secretion system (11). The secretion system exports subunits to polymerize the axle-like rod through the peptidoglycan and the outer membrane (if present), and beyond that a short, extracellular curved structure called the hook (12). Hook completion changes the specificity of the secretion system (13–15) to activate the alternative sigma factor SigD and late class flagellar filament proteins, by exporting the anti-sigma factor FlgM (16–19). Assembly of the 8-13 nm wide rod (20–24) through the envelope is problematic, as its width is seemingly incompatible with the 4 nm average pore size of peptidoglycan (2). Some Gram negative bacteria are thought to mount peptidoglycan lyases on the end of the extending rod to remodel the thin layer of PG at the point of contact and make room for rod transit (25–28). Gram positive bacteria however, have thick multi-lamellar peptidoglycan and how the flagellum transits the Gram positive cell envelope is unknown.

*Bacillus subtilis* is a Gram-positive oblong bacterium that synthesizes approximately 20 flagella per cell distributed in a non-random grid-like pattern along its length (29). While *B. subtilis* encodes over 40 putative peptidoglycan lyase proteins, neither reverse nor forward genetic approaches have been able to identify a lyase specific for flagellar assembly (30–33). Recent work indicated that flagellar basal bodies of *B. subtilis* are mobile in the membrane early in assembly and become immobilized once the rod/hook structure that transits the envelope is complete (34). A model was proposed suggesting that the porosity of peptidoglycan was heterogenous, and that nascent basal bodies diffuse in the membrane until encountering a hole of sufficient diameter that permits rod assembly. The model also suggested that large pores might be non-randomly distributed across the peptidoglycan thereby serving as a template for grid-like flagellar patterning. In short, rather than using a lyase to make holes during synthesis, it was proposed that nascent flagella in *B. subtilis* randomly search for permissive sites pre-existent in the peptidoglycan cell wall.

Peptidoglycan porosity is thought to be governed by the frequency of amino acid crosslinks between the polysaccharide chains mediated in part by extracellular penicillin binding proteins (PBPs) (4,35,36). *B. subtilis* encodes a number of PBPs (37,38), but evidence suggests that crosslinking by PBP1 in particular plays a role in decreasing the frequency and/or diameter of pores in the PG mesh (39–42). Here we explore the relationship between flagella assembly and the porosity of peptidoglycan by genetically mutating PBP1. We find that mutation of PBP1 leads to a loss of flagellar-mediated swarming motility, but also confers a severe defect in growth rate. The growth rate defect was dependent on flagellar synthesis, and induction of flagellar synthesis genes lead to cell lysis. Further characterization indicated that the requirement for flagellar synthesis was due to activation of the alternative flagellar sigma factor SigD and the multiple peptidoglycan hydrolases under its control. Only a subpopulation of cells activates SigD in *B. subtilis* (43,44), and the toxicity of the flagellum in those cells is counteracted by the activity of PBP1 and the stress response regulator Spx (45,46). In sum, our data suggest that flagellar biosynthesis can have detrimental effects on cell viability and that the cell encodes multiple systems to neutralize its toxicity.

## RESULTS

### Flagellar synthesis is toxic in the absence of PBP1

To test the effect of alterations in peptidoglycan porosity on flagellar basal body pattern, the *ponA* gene encoding PBP1, was mutated in the ancestral strain of *B. subtilis*, NCIB3610 (hereafter 3610). While the *ponA* mutant was defective for swarming motility (**Fig S1**), cells nonetheless produced puncta of flagellar basal bodies as indicated by a green fluorescent protein fusion to the cytoplasmic flagellar structural protein FliM (29,34) (**Fig. 1**). Puncta pattern analysis was complicated however, as the *ponA* mutant appeared to produce fewer but more intense FliM-GFP foci than the wild type and the foci also appeared to be less evenly distributed across the cell length (**Fig. 1**). The large foci of fluorescence likely indicates the aggregation of multiple basal bodies in the membrane (29), and the cells themselves were frequently misshapen, elongated, and often appeared to have undergone lysis. Perhaps consistent with a high frequency of death, the *ponA* mutant also exhibited a severe defect in growth rate (**Fig. 2A**). The swarming, basal body distribution, and viability phenotypes were due to the absence of PBP1 as each phenotype was complemented when the *ponA* gene was expressed from its native promoter and inserted at an ectopic site in the chromosome (**Fig 1**, **Fig 2A, Fig S1**). We conclude that flagellar motility and patterning were perturbed in the absence of PBP1, but the defect in growth was the most prominent phenotype, and one that complicated the interpretation of the others.

**Figure legend 1.**
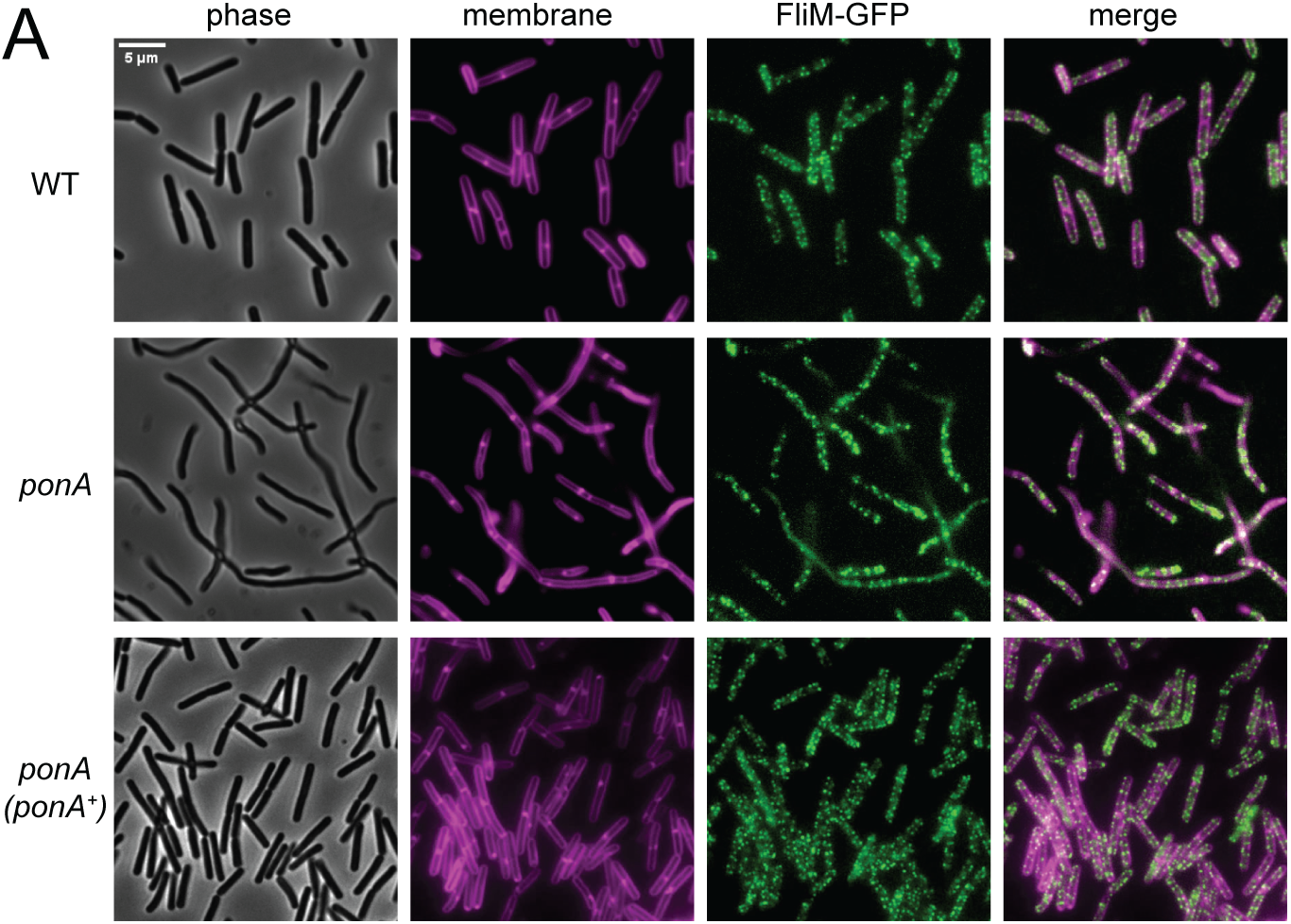
Cells mutated for PBP1 are defective in flagellar basal body synthesis and cell morphology. Micrograph images of phase contrast and fluorescence microscopy of membranes stained with FM4-64 (false colored magenta), flagellar basal bodies marked with FliM-GFP (false colored green), a merge of the two fluorescence channels. The *trans* complementation construct in which *ponA* was cloned under its native promoter and inserted at an ectopic site in the chromosome *ponA (ponA^+^*). The following strains were used to generate the figure: WT (DK1906), *ponA* (DB2642), and *ponA* (*ponA^+^*) (DB3420). Scale bar is 5 µm and is the same for all panels.

**Figure legend 2.**
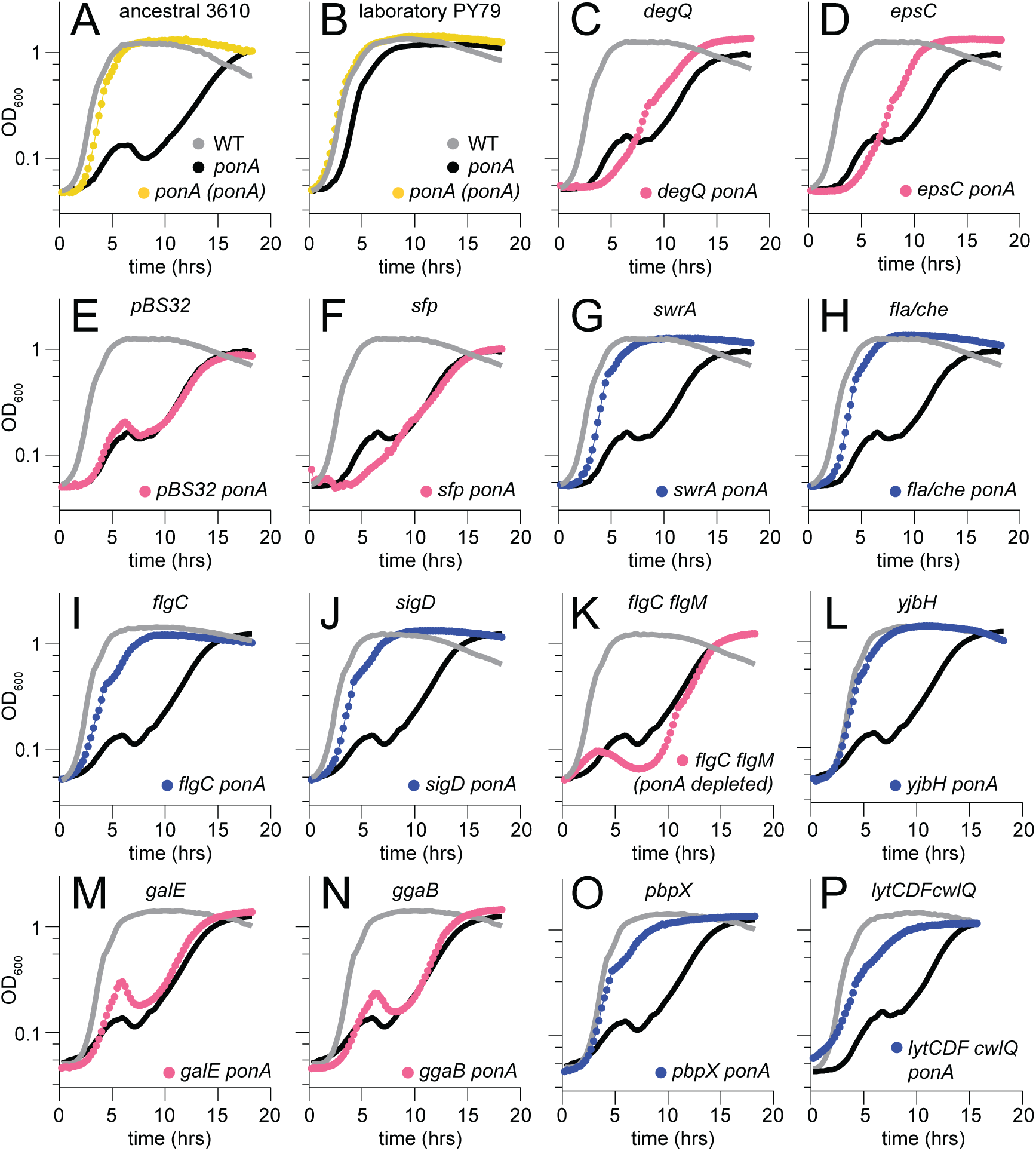
Cells mutated for PBP1 have a growth defect that can be rescued by the inactivation of SigD, YjbH, and PbpX. Each graph plots the optical density measured over time of cells grown in microtiter dishes with LB broth, shaken at 37°C. Gray lines indicate the wild type and black lines indicate the *ponA* mutant. All graphs use the ancestral 3610 genetic background except for **panel B** which uses the laboratory strain PY79 genetic background. **Panel K** involves a *ponA flgC flgM* triple mutant with a copy of *ponA* expressed from the IPTG-inducible *P_spank_* promoter for PBP1 depletion in the absence of IPTG, as combinations of *ponA* and *flgM* mutations were genetically unobtainable. Lines in yellow indicate *trans* complementation constructs in which *ponA* was cloned under its native promoter and inserted at an ectopic site in the chromosome (“*ponA (ponA)*”). Pink and blue lines indicate cells doubly mutated for *ponA* and the gene indicated. Pink lines indicate mutations that failed to restore robust growth to the *ponA* mutant while blue lines indicate mutations that restored growth to near wild type levels. The following strains were used to generate the panels: **A)** 3610 WT (DK1042), 3610 *ponA* (DB2291), 3610 *ponA (ponA)* (DB2699), **B)** PY79 WT (PY79), PY79 *ponA* (DB2329), PY79 *ponA (ponA)* (DB2699), **C)** *degQ ponA* (DB2384), **D)** *epsC ponA* (DB2552), **E)** *pBS32 ponA* (DB2558), **F)** *sfp ponA* (DB2381), **G)** *swrA ponA* (DB2383), **H)** *fla/che ponA* (DB2382), **I)** *flgC ponA* (DB2274), **J)** *sigD ponA* (DB2493), **K)** *flgC flgM* (*ponA* depleted) grown in the absence of IPTG (DB2643), **L)** *yjbH ponA* (DB2858), **M)** *galE ponA* (DB2859), **N)** *ggaB ponA* (DB2829), **O)** *pbpX ponA* (DB2869). Each data point is the average of three replicates.

The growth defect in the absence of PBP1 was severe in the ancestral strain genetic background, but growth defects have not previously been reported for *ponA* mutants in laboratory strains under similar conditions (42,47–49). Indeed, the same *ponA* allele introduced to laboratory strain PY79 did not confer a dramatic growth defect suggesting that mutations unlinked to *ponA* were responsible for the growth differences (**Fig 2B**). As laboratory strains like PY79 are genetic descendants of 3610 (50), *ponA* was mutated in a 3610 genetic background that was also mutated in genes known to be disrupted by laboratory strain polymorphisms (51–55). Mutation of the *epsC* gene, *degQ* gene, *sfp* gene, or elimination of the 86 kb plasmid pBS32 failed to substantially enhance growth of the *ponA* mutant (**Fig 2C-F**). By contrast, mutation of the *swrA* gene restored growth of the *ponA* mutant to levels resembling that of the laboratory strain (**Fig 2G**). We conclude that the loss-of-function frameshift mutation in *swrA* in laboratory strains permits rapid growth of the *ponA* mutant. Conversely, we conclude that in the ancestral background, the functional allele of *swrA* is toxic when PBP1 is inactivated.

The *swrA* gene encodes SwrA, a small protein which activates the 27 kb *fla/che* operon dedicated to flagellar biosynthesis (56–58). Like mutation of *swrA*, deletion of the entire *fla/che* operon also rescued growth in the absence of PBP1 (**Fig 2H**). To determine whether activation of the *fla/che* operon was sufficient to cause death, the SwrA-enhanced *P_fla/che_* promoter was replaced with a promoter that was IPTG-inducible (29). Cells of a *ponA* mutant in which the *P_fla/che_* promoter had been replaced grew rapidly in the absence of IPTG, but growth was abolished when 1 mM IPTG was present. By varying inducer concentration, the toxicity of *fla/che* operon expression was titrated to a narrow three-fold dynamic range (**Fig 3A**) and one hour after induction, extensive cell lysis was observed (**Fig 3B**). We conclude that activation of the *fla/che* operon in the absence of PBP1 causes growth defects and cell lysis.

**Figure Legend 3.**
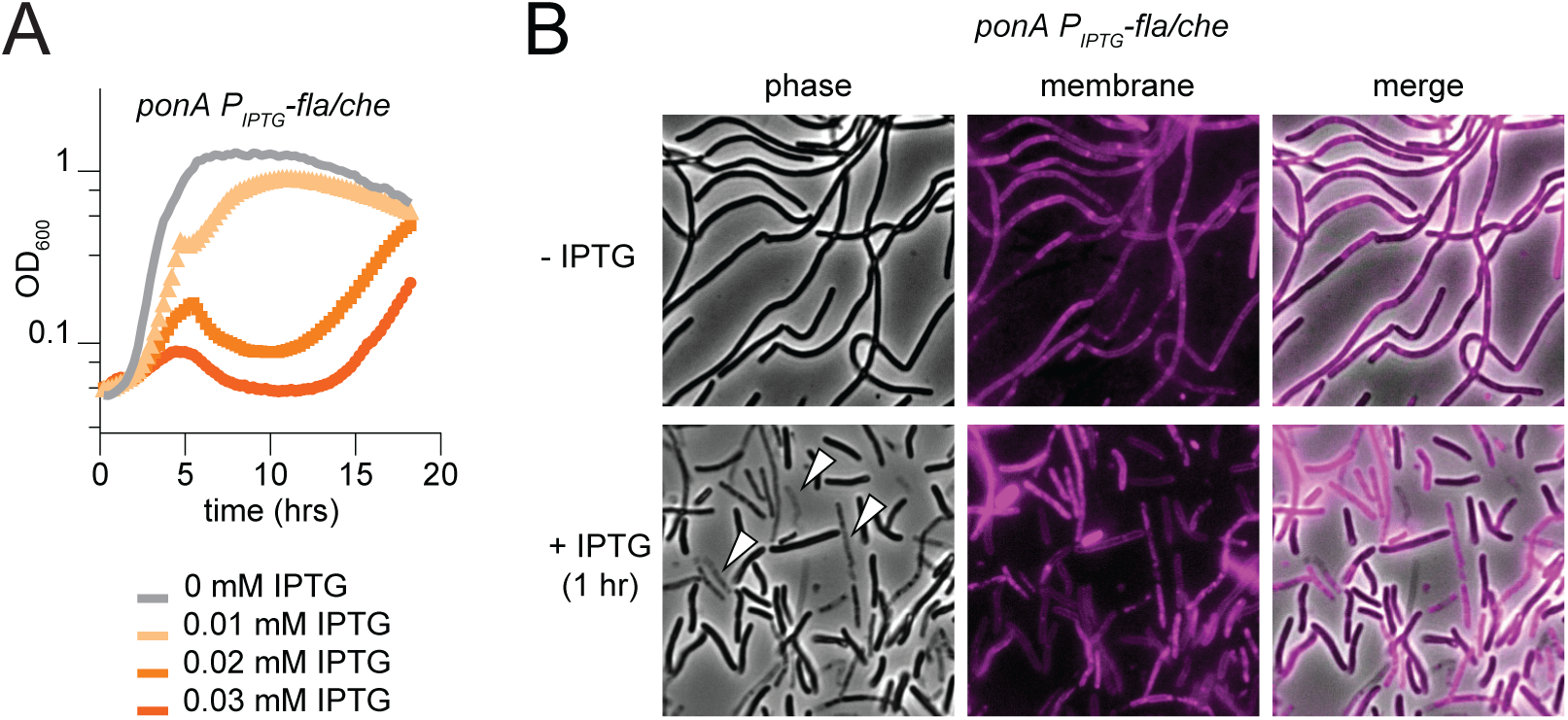
Induction of the fla/che operon or overexpression of PBPX is lethal in the absence of PBP1. Each graph plots the optical density measured over time of cells grown in microtiter dishes with LB broth, shaken at 37°C. Panel A) Growth curves of a strain mutated for ponA and in which the fla/che operon was placed under the control of the IPTG-inducible *P_hypsank_*promoter. The gray line indicates growth in the absence of IPTG while the increasing intensity of orange lines indicate growth in the presence of increasing amounts of IPTG, as indicated. The following strain was used for all lines in the panel: *P_hyspank_-fla/che ponA* (DB2587). Panel B) Induction of the fla/che operon causes lysis in the absence of PBP1. Phase contrast and fluorescence microscopy (membranes stained with FM4-64, false colored red), in the absence (- IPTG) and growth for one hour in the presence of 1 mM IPTG (+ IPTG).

### Flagellar toxicity in the absence of PBP1 requires the activation of SigD, and the presence of PbpX

The *fla/che* operon encodes 32 genes dedicated to the synthesis of early stages of the flagellar structure (*fla*) and the chemotactic control of motile behavior (*che*) (59–62). To test the role of the flagellar structural genes, cells were mutated for *flgC*, encoding the rod structural protein FlgC thought to be the first component of the flagellar rod to enter the peptidoglycan layer (34,63) (**Fig 4A**). Cells mutated for FlgC restored robust growth in the absence of PonA (**Fig 2I**). The absence of FlgC may have restored growth either due to the structural absence of the flagellar rod, or for regulatory reasons, as completion of the rod, and later the hook, is necessary for activation of the alternative sigma factor SigD (19,64). Consistent with a regulatory role for FlgC, mutation of the gene encoding SigD, found at the 3’ end of the *fla/che* operon, also enhanced growth in the absence of PonA (**Fig 2J**) (61,65). Furthermore, poor growth was restored with SigD activity in cells doubly mutated for FlgC and the SigD-antagonist FlgM, upon *ponA* depletion (**Fig 2K**). We conclude that expression of the *fla/che* operon is toxic in the absence of PBP1 because completion of the flagellar rod-hook structure results in the activation of SigD.

**Figure Legend 4.**
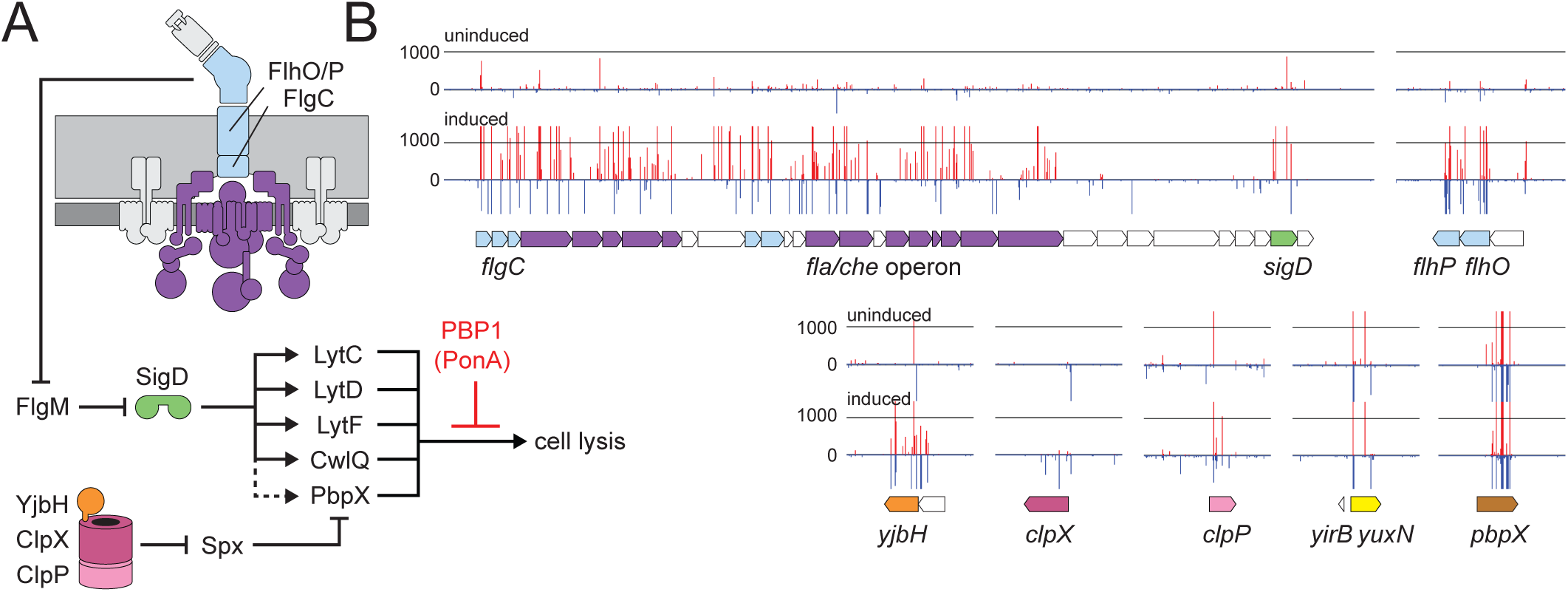
Mutation of genes encoding early flagellar structural proteins, SigD, proteins that inhibit Spx accumulation and PBPX improve growth in the absence of PBP1. Panel A) Cartoon model of flagellar structure and genetic wiring diagram of flagellar toxicity. Top, dark gray bar indicates plasma membrane and medium gray bar indicates peptidoglycan. Purple shading indicates components of the flagellar type III secretion system, flagellar basal body, and flagellar C-ring, while light blue shading indicates components of the flagellar rod and hook. Flagellar hook completion triggers export of the anti-sigma factor FlgM which relieves inhibition on the flagellar sigma factor SigD (green). SigD in turn directly activates a regulon of genes including four peptidoglycan lyases (LytC, LytD, LytF, and CwlQ), and either directly or indirectly activates the expression of PbpX which combine promote cell lysis. The lytic activity of the PbpX-enhanced peptidoglycan lyases is antagonized by PBP1 (red) and Spx. Spx levels are restricted by regulatory proteolysis governed by the ClpX/ClpP cytoplasmic protease and the specificity adaptor YjbH. Panel B) Transposon sequencing data from strain DB2587 mutated for *ponA* and in which the *fla/che* operon was placed under the control of the IPTG-inducible *P_hypsank_* promoter, and mutagenized with the transposon TnWX642. Approximately 1 million transposant colonies were separately harvested in the absence (top graphs) and presence (bottom graphs) of IPTG and bulk insertions from both pools were sequenced Illumina sequencing. Bars indicate the locations of insertions with respect to gene position indicated below. The length of the bars indicates the number of insertions at that position with a black line indicating 1000 insertions at that position. Red bars indicate insertions of the transposon in one orientation while blue bars indicate insertions the other. Each genetic map below indicates gene orientation as arrows and all regions are expressed at the same scale. Separations between the data indicate unlinked loci. For the sake of simplicity, neighboring genes have been omitted unless part of an operon. Note that mutations that improved growth of PBP1 hyperproliferated in both the absence and presence of IPTG so peak loss is uninterpretable while peak enhancement is exaggerated.

SigD directs RNA polymerase to express a regulon of genes (43,66,67), but a reverse genetic approach in which 21 out of 23 SigD-activated genes were separately mutated failed to enhance growth in the absence of PBP1 (**Fig S2**). The remaining two mutants restored growth but were defective in FlhO and FlhP, structural subunits of the distal rod which while under partial SigD control, are also required for SigD activity (63,68) (**Fig 4A**). We next undertook an unbiased transposon mutagenesis and sequencing approach (TnSeq) to determine which genes, under SigD-control or otherwise, were required for growth inhibition when the *fla/che* operon was expressed in the absence of PonA. Briefly, a PBP1 mutant strain with the IPTG-inducible *fla/che* operon was transposon mutagenized in both the presence and absence of inducer, DNA was extracted from both pools, sequenced, and compared. We found that the transposon insertion diversity was low for both pools due to the proliferation of mutants that improved growth and/or survival without PBP1 in general. Nonetheless, the transposon insertion frequency within the *fla/che* operon dramatically increased when inducer was present, supporting previous observations that induction of the operon was toxic (**Fig 4B**). Ultimately, no genes known to be SigD regulated, besides *flhO* and *flhP* (**Fig 4B, Fig S3A**), were found to experience an increase in insertion density when the *fla/che* operon was induced in the absence of PBP1.

We considered the possibility that there might be previously unrecognized members of the SigD-regulon responsible for toxicity, and three genes with high insertion frequency were tested individually: *galE*, encoding UDP-glucose epimerase (69), *ggaB*, encoding a protein required for an N-acetylgalactosamine-containing wall teichoic acid (70), and *pbpX*, encoding a putative class C penicillin binding protein homolog of unknown function (**Fig S3B**). While mutation of *galE*, or *ggaB*, had no effect on growth in any background (**Fig S4A**), mutation of *pbpX,* restored robust growth in the absence of PBP1 (**Fig 2O**). Next, an artificial expression system was generated in which expression of *pbpX* was placed under the IPTG-inducible *P_hyspank_*promoter and inserted at an ectopic location in the chromosome. Artificial induction of PbpX inhibited growth in the absence of PBP1 but had no phenotype when expressed in wild type cells (**Fig 5**). We conclude that the expression of PbpX is necessary and sufficient to impair cell viability when PBP1 is absent.

**Figure Legend 5.**
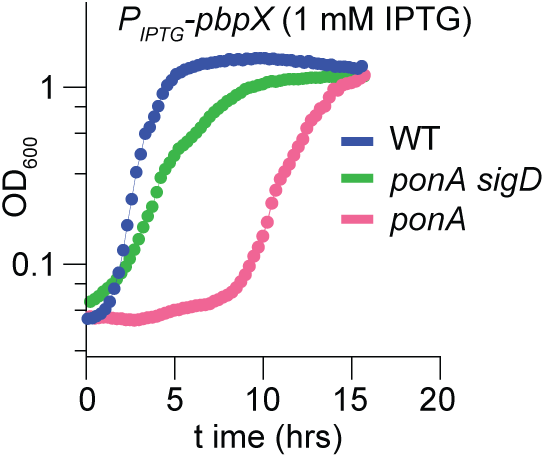
PBPX expression is activated by SigD and repressed by Spx. Each graph shows the transcriptional activity of *P_pbpX_* in various genetic backgrounds by measuring β-galactosidase activity of a *P_pbpX_-lacZ* transcriptional reporter fusion. Each bar shows the mean Miller units of three biological replicates, with error bars indicating standard deviation. The following strains were used to generate these panels: **A)** WT (DB3109), *sigD* (DB3135), *flgM* (DB3136), *sigD flgM* (DB3608). **B)** WT (DB3109), *yjbH* (DB3193), *spx* (DB3605)*, yjbH spx* (DB3194), **C)** *sigD yjbH* (DB3606)*, sigD spx* (DB3575)*, yjbH flgM* (DB3576).

To determine whether *pbpX* gene expression was under SigD control, a transcriptional reporter was generated by fusing the promoter region of *pbpX* (*P_pbpX_*) upstream of the *lacZ* gene, encoding β-galactosidase, and inserted at an ectopic location in various genetic backgrounds. The *pbpX* gene is not known to be a direct target of SigD, but appeared to be at least partially SigD-activated, as mutation of SigD reduced its expression (**Fig 6A**). Mutation of the SigD antagonist FlgM however, did not result in increased *pbpX* expression, as is often the case for genes under SigD-control, and simultaneous mutation SigD and FlgM gave inconclusive epistasis (**Fig 6A**). Due to dissimilar behavior with other members of the SigD regulon, we suspect that SigD activation of *pbpX* is likely indirect, and we note that *pbpX* experienced a dramatic enrichment in TnSeq insertion density in the absence of PBP1 whether or not the flagellar genes were induced (**Fig 4B**). Moreover, over-expression of *pbpX* in the absence of both PBP1 and SigD failed to fully inhibit growth suggesting that *pbpX* expression was no longer sufficient to cause toxicity (**Fig 5**). Whether or not *pbpX* is either directly or indirectly activated by SigD, we conclude that flagellar toxicity requires both PbpX and one or more as-yet-unidentified genes under SigD-control.

**Figure Legend 6.**
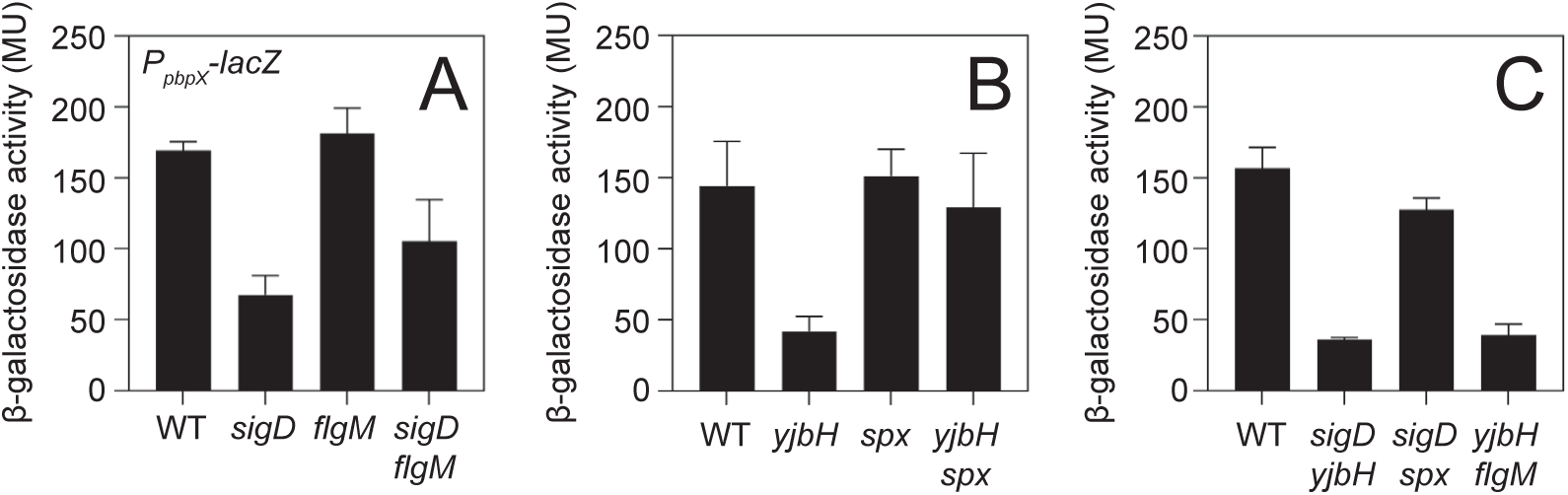
Spx accumulates in the absence of PBP1 when the *fla/che* operon is induced. Western blot analysis was performed on cell lysates extracted from the genetic backgrounds indicated above. The Western blots were developed using a primary antibodies raised against Spx and primary antibodies raised against the vegetative sigma factor SigA to serve as a loading control. Strains containing the IPTG-inducible fla/che operon (*P_hyspank_-fla/che*) were grown in the presence (+) and absence (–) of IPTG. The following strains were used to generate these data: WT (DK1042), *yjbH* (DS8875), *ponA* (DB2291), *ponA P_hyspank_-fla/che* (DB2279), *flgM* (DS322)

### SigD toxicity is due to the expression of multiple peptidoglycan lyases and is overcome by high levels of Spx

To learn more about the mechanism of flagellar toxicity, we focused on genes that displayed an increase in insertion frequency that were related to the stress response factor Spx (71–72) (**Fig S3C**). For example, when the *fla/che* operon was induced, there was an increase in the insertion frequency in *yjbH*, encoding the adaptor that delivers Spx to the ClpXP protease for degradation (73–75), and insertion frequency also increased for both genes encoding the protease subunits (76) (**Fig 4A, 4B**). Moreover, insertion frequency was high in both the induced and uninduced conditions for *yuxN*, encoding a DNA binding transcription factor homolog that represses YirB, a small anti-YjbH anti-adaptor protein that promotes Spx accumulation (77,78) (**Fig 4B**). Mutation of YjbH restored robust growth in the absence of PBP1 (**Fig S4B**) and combined, the gene set suggested that high levels of Spx could ameliorate the toxicity of SigD activity, either prior to or during flagellar activation. Spx is a monomeric transcription factor that interacts with RNA polymerase to activate and repress a wide range of genes related to surviving cellular stress (72,79–81).

To determine whether Spx levels rose when flagella were synthesized in the absence of PBP1, Western blot analysis was conducted on a variety of strain backgrounds using primary antibodies to Spx, and the vegetative sigma factor SigA as a loading control. Consistent with previous reports, Spx protein was barely detectable in wild type cells, and Spx accumulated to high levels when the proteolytic adaptor protein YjbH was disrupted (**Fig 7**) (73,74). Mutation of PBP1 appeared to cause a slight increase in Spx relative to wild type (**Fig 7**). Moreover, Spx levels were correlated with the expression of the *fla/che* operon as Spx decreased in the absence of PBP1 when the *fla/che* operon was uninduced and increased after 1 hour of *fla/che* operon induction, a timepoint concurrent with widespread lysis (**Fig 7**). Elevation of SigD activity alone by mutation of FlgM also appeared to cause a slight increase in Spx even when PBP1 was present (**Fig 7**). We conclude that *fla/che* operon expression and SigD activity causes cellular stress and that either the presence of PBP1 and/or the accumulation of Spx could ameliorate toxicity.

**Figure 7.**
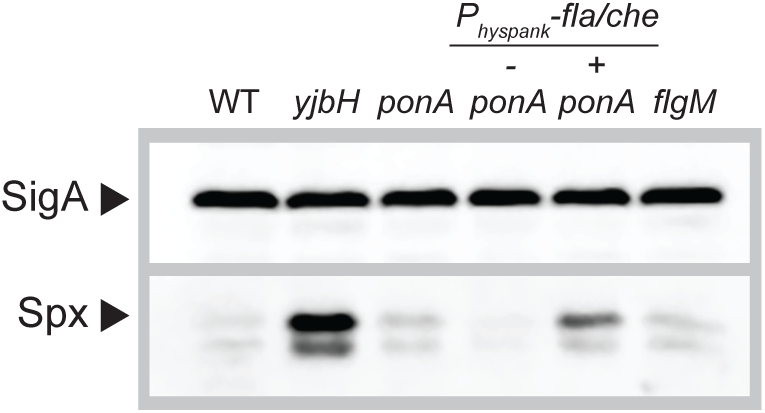

One way in which Spx might prevent SigD-dependent toxicity is by repressing expression of PbpX. To test the role of Spx in PbpX regulation, the *P_pbpX_-lacZ* reporter was measured in a variety of genetic backgrounds. Consistent with Spx-mediated inhibition, PbpX expression decreased in cells mutated for YjbH (**Fig 6B**). Moreover, while mutation of Spx alone had no effect on *P_pbpX_*, simultaneous mutation of both Spx and YjbH indicated that mutation of Spx was epistatic (**Fig 6B**). Next we sought to determine the relationship between SigD-mediated activation and Spx-mediated inhibition of *P_pbpX_*. Simultaneous mutation of both SigD and YjbH did not appear to have an additive effect on expression, suggesting perhaps that they were acting through a similar intermediate (**Fig 6C**). Furthermore, simultaneous mutation of both SigD and Spx elevated PbpX expression such that it resembled the Spx mutation alone, while simultaneous mutation of YjbH and FlgM, resembled mutation of YjbH alone (**Fig 6C**). We conclude that the epistasis analysis supports the idea that SigD activates PbpX indirectly, and that SigD activity creates a general stress response that can be overridden by high levels of Spx, at least at the level of PbpX gene expression.

A number of observations indicated that SigD-toxicity was due to cell wall-related stress. First, induction of the *fla/che* operon in the absence of PBP1 appeared to cause cellular lysis, and second, high levels of Spx was protective and has been shown enhance the resistance of *B. subtilis* to cell wall damage (45,78). Third, mutation of PbpX was found to improve the growth of cells mutated for PonA, and PbpX is homologous to a group of small class 3 penicillin binding proteins that includes FmtA from *Staphylococcus aureus* (82) (**Fig S5**). FmtA has been shown to remove D-alanine groups from Gram-positive teichoic acid (83), and teichoic acid alanylation has been associated with the control of autolytic peptidoglycan lyases (84–87). SigD directly activates at least four peptidoglycan lyases including LytC, LytD, LytF, and CwlQ (31,88–90) and we wondered whether lyase synergy might explain why no single gene known to be under SigD control appeared necessary for toxicity in the absence of PBP1 during TnSeq analysis. Indeed, cells simultaneously mutated for LytC, LytD, LytF and CwlQ restored growth in liquid media in the absence of PBP1 (**Fig 2P**), and resembled cells mutated for PBP1 and PbpX (**Fig 2O**). We conclude that toxicity in the absence of PBP1 requires flagellar hook completion, which in turn activates SigD, which in turn activates a suite of peptidoglycan lyases, the activity of which is enhanced by PbpX and antagonized by PBP1 (**Fig 4A**).

## DISCUSSION

Flagella are complex, multi-subunit molecular machines that transit all layers of the bacterial cell envelope, but flagella are wider than the average pore size of peptidoglycan. In Gram negative bacteria, lyases are thought to create holes for the axle-like rod, but no dedicated lyase necessary for flagellar assembly has been discovered in Gram positive bacteria. Instead, a recent model has suggested that Gram positive flagellar basal bodies diffuse in the membrane and are captured at periodic pre-existent holes in the peptidoglycan of sufficient diameter (34). In an effort to increase the frequency of flagellar permissive sites we mutated PBP1, a protein previously shown to reduce peptidoglycan porosity, and found that cells exhibited a high rate of lysis due to flagellar assembly itself. Subsequent genetic analysis indicated that toxicity was due to completion of the flagellar hook and activation of the alternative flagellar sigma factor SigD, an event that occurs after peptidoglycan transit. Once activated, SigD was toxic due to parallel expression of multiple peptidoglycan lyases, which appear to cause extensive cell wall damage and autolysis when PBP1 is absent. Thus, our results indicate that flagellar assembly, not normally associated with cell viability, can under certain conditions have toxic side effects.

Flagellar toxicity is minimized by PBP1, a member of the Class A penicillin binding proteins that have both transglycosylase activity to extend peptidoglycan polysaccharide chains, as well as transpeptidase activity to crosslink the side chain amino acids (35,91,92). As such, PBP1 was once thought to be central to cell wall synthesis, but mutation of PBP1 in *B. subtilis* laboratory strains had little phenotypic effect on growth, a fact that would later lead to the discovery that the SEDS proteins were the primary peptidoglycan synthases in bacteria (93).

Thus, the study of cell wall synthesis in laboratory strains that are naturally reduced for flagellar synthesis was a fortunate happenstance, as otherwise, the absence of PBP1 might have appeared to be misleadingly severe. We find that the strain dependent difference is due to the ancestral strain of *B. subtilis* being functional for SwrA, an activator of the *fla/che* operon that increases the number of flagella synthesized per cell and increases the frequency of cells with flagella (43,58). We suggest that laboratory strains defective for SwrA have greater tolerance for mutation of PBP1 due to the high frequency of cells that are in the SigD-OFF state (43,44). Whatever the case, we find that mutation of PBP1 is conditionally essential based on the level of flagella expression (**Fig 3A**).

Current models suggest that the function of PBP1 is to repair holes in the peptidoglycan superstructure. As evidence for repair, the class A PBPs are not required for cellular elongation (37,39,47,93) and instead increase integrity and prevent cell lysis (48,94,95). Moreover, PBP1 localizes and moves independently of the SEDS-containing cellular elongation complex (39,40), and PBP1 movement is arrested, presumably during activity, in response to the presence of PG damaging agents or limiting PG precursors (39,41). Finally, PBP1 of *B. subtilis* has a long, intrinsically disordered, C-terminal domain proposed to sense and localize to holes in the Gram-positive wall superstructure (42). Here we find that flagellar synthesis triggers a cell wall damage response thereby providing a venue for PBP1-directed repair (**Fig 7**, 45,46,78). Thus, PBP1 activity likely supports growth either by blocking the activity of a suite of flagellar-regulated peptidoglycan hydrolases or it repairs the damage faster than it can accumulate.

Transposon sequencing identified two other factors related to cell wall damage besides the flagellar structural genes, which when mutated, improve growth in the absence of PBP1. First, growth was improved by disrupting the regulatory proteolysis system that keeps Spx levels low in the cytoplasm (81). Spx is a monomeric regulator that binds to RNA polymerase and has pleiotropic effects on cell physiology, including the activation of multiple regulons dedicated to cell wall damage repair (45,96). The second factor is PbpX, a poorly-studied Class C penicillin binding protein homolog, which improved growth in the absence of PBP1 whether or not the flagellar genes were expressed, and we note that the only other report of a *pbpX* mutant phenotype was in a screen for the improved growth in the presence of the PBP1 inhibiting β-lactam antibiotic cefuroxime (96). The closest PbpX relative of known function is FmtA of *Staphylococcus aureus,* an enzyme that removes D-alanine modifications from teichoic acid (**Fig S5**; 82,83). Teichoic acids are important for the binding and activity of peptidoglycan lyase enzymes in general, and for the SigD-regulated lyases in particular (84–86,97–100). If PbpX functions like FmtA, we infer that D-alanylation of teichoic acids inhibits or otherwise restricts the activity of the peptidoglycan lyases under flagellar control.

*B. subtilis* encodes more than 30 proteins that are known or predicted to cleave peptidoglycan at different bonds in the superstructure (33), and four of the lyases are under SigD control. Two of the SigD-dependent lyases appear to directly support motility, as the muropeptidase LytF promotes cell separation after division to yield motile individuals (30, 101,102) and the lytic transglycosylase CwlQ promotes swarming by increasing flagellar torque (31,103). The remaining two lyases include the muramidase LytC that is thought to be the primary enzyme involved in cell wall turnover and autolysis (84,104–106) and the glucosaminidase of unknown function LytD (30,89,105), which together appear to indirectly promote hyperflagellation during swarming (30,107). That four diverse peptidoglycan lyases are expressed under flagellar control in *B. subtilis* may speak to the complexities of managing a transenvelope nanomachine in the context of the Gram positive cell architecture. None of the lyases are required for flagellar transit either singly or together, and a combination of the lyases were needed for toxicity.

Flagellar synthesis is highly regulated at multiple levels, and regulation is considered important to restrict the high metabolic cost of the subunits needed for assembly. Here we show that flagellar synthesis in *B. subtilis* comes at an additional cost of cell wall integrity that is ameliorated by PBP1 and exacerbated by PbpX. While most dramatic in the absence of PBP1, flagellar toxicity may be subtly pervasive even in wild type as we note that a frameshift mutation in SwrA appears in the earliest *B. subtilis* laboratory strains, perhaps selected to promote growth by relegating flagellar-mediated cell wall stress to a minority subpopulation (43,50,108). If true, domestication to laboratory conditions may have reduced flagella in bacteria besides *B. subtilis*, perhaps obscuring their conditionally toxic effect, or perhaps flagellar toxicity has simply gone unnoticed because cell wall synthesis and motility are separate fields and seldom studied in tandem (109,110). Whatever the case, pursuit of an alternative model of flagellar insertion and grid-like patterning lead to the discovery that flagellar synthesis can have toxic side effects. We note that flagella are but one example of a large complex that transits the peptidoglycan by poorly understood mechanisms, and we wonder whether cell wall integrity may require special management during the synthesis of other motility machines and secretion apparati.

## MATERIALS AND METHODS

### Strains and growth conditions

*B. subtilis* strains were grown in Luria-Bertani (LB) (10 g tryptone, 5 g yeast extract, 5 g NaCl per L) broth or on LB plates fortified with 1.5% Bacto agar at 37°C. When appropriate, antibiotics were included at the following concentrations: 10 μg/ml tetracycline, 100 μg/ml spectinomycin, 5 μg/ml chloramphenicol, 5 μg/ml kanamycin, and 1 μg/ml erythromycin plus 25 μg/ml lincomycin (*mls*). Isopropyl b-D-thiogalactopyranoside (IPTG, Sigma) was added to the medium at the indicated concentration when appropriate.

### Microscopy

Fluorescence microscopy was performed on a Nikon Ti2E microscope equipped with Plan Apo 100x/1.4NA phase contrast oil objective and a Prime 95B camera. Strains of *B. subtilis* were grown in LB (with antibiotics or IPTG as indicated) at 37c to an exponential OD_600_ (0.4-0.8). To visualize membranes, cells were pelleted and resuspended in 30 μL of 1x PBS containing 15 μg/ml of FM4-64 (Invitrogen #T13320), incubated for 2 minutes at room temperature in dark and washed before imaging. To visualize flagellar filaments, cells were pelleted and resuspended in 50 μL of 1× PBS containing 5 μg/mL Alexa Fluor 488 C_5_ maleimide (Invitrogen; molecular probes A10254) incubated for 3 minutes in the dark at room temperature. Cells were subsequently washed with 1mL 1x PBS and pelleted again. The pellets were then resuspended in 30ul of 15 μg/ml of FM4-64, incubated for 2 minutes in the dark at room temperature, washed and resuspended in 30 μL 1x PBS. Cells were immobilized using 1% agarose pads made with water and images were captured with NIS-Elements software. Images were cropped and adjusted using ImageJ software.

### Growth Curves

*B. subtilis* strains were inoculated into 3mL LB media, grown at 37°C in a roller drum until cultures were turbid, and back diluted to an OD600 of 0.01. 500uL of culture was then loaded in technical triplicate into a Greiner Cellstar 48 well plate with no lid. Strains expressing the *P_hyspank_-fla/che operon* construct were initially grown without IPTG, and indicated concentrations of IPTG were added at the start of the growth curve. Strains expressing *P_spank_-ponA* were grown with IPTG, then washed and resuspended in either plain LB or LB supplemented with 1 mM IPTG as indicated at the start of the growth curve. 48 well plates were loaded onto a BioTek Synergy H1 microplate reader and growth curves were measured using BioTex microplate reader and imager Gen5 3.11 software via the following protocol. Procedure was set as a “kinetic run” for 18 h with absorbance readings of 600 A every 15 min. The plate was set to shake in a continuous double-orbital at 37°C.

### Western blotting

*B. subtilis* strains were grown in LB to OD_600_ ∼1.0, 1 ml was harvested by centrifugation, and resuspended to 10 OD_600_ in Lysis buffer (20 mM Tris pH 7.0, 10 mM EDTA, 1 mg/ml lysozyme, 10 mg/ml DNAse I, 100 mg/ml RNAse I, 1 mM PMSF) and incubated 30 minutes at 37°C. 10 ml of lysate was mixed with 2 ml 6x SDS loading dye. Samples were separated by 12% Sodium dodecyl sulfate-polyacrylamide gel electrophoresis (SDS-PAGE). The proteins were electroblotted onto nitrocellulose, probed with either a 1:2,000 dilution of anti-Spx primary antibody or 1:80,000 dilution anti-SigA primary antibody, and developed with 1:10,000 dilution of horseradish peroxidase-conjugated goat anti-rabbit immunoglobulin G secondary antibody and Immun-Star HRP developer kit (Bio-Rad).

### TnSeq

We performed TnSeq using the *mariner* transposon delivery plasmid pWX642 (111). This plasmid contains (i) an erythromycin resistance gene for plasmid selection (ii) a spectinomycin resistance gene flanked by inverted repeats that are recognizable by the transposase (iii) an MmeI site inside one of the inverted repeats to facilitate DNA library prep and determine the orientation of Tn insertion, (iv) the HiMar transposase under constitutive expression to allow Tn hopping, and (v) a temperature-sensitive replication origin for *B. subtilis*, which allows plasmid loss at high temperature. After plasmid loss, the spec resistant colonies contain a permanent chromosomal Tn insertion that is no longer mobile due to the loss of transposase. To generate the strains for mutagenesis, DB2587 (*P_hyspank_-fla/che operon kan ponA::tet*) was transduced with lysate from DK6865 (carrying the pWX642 plasmid), plated on LB agar containing mls and kan (to maintain *P_hyspank_-flache operon kan*), and incubated at 30°C for 48 h. Four individual colonies from the transductants were inoculated into one tube containing 4 mL LB broth supplemented with spec and kan and incubated at 22°C for 20 h, and this growth procedure was repeated with 4 parallel tubes, which were eventually pooled together. To generate the mutant libraries, the mutagenized pools were diluted appropriately, plated on 20 large (150 by 15 mm) LB spec kan agar plates, +/-IPTG, and incubated at 42°C overnight such that each plate had ∼50,000 colonies. Colonies were harvested with a rubber scraper, the slurry was diluted to an OD_600_ of 5, and genomic DNA was purified using the DNeasy blood and tissue DNA purification kit (Qiagen).

Next, 3 ug of gDNA was digested with MmeI for 90 min, and subsequently treated with quick calf intestinal alkaline phosphatase (CIP) for 60 min at 37°C. The digested DNA was extracted using phenol-chloroform, precipitated using ethanol, and resuspended in 15 μL nuclease free ddH2O. Subsequently, the digested DNA end was ligated to an annealed adapter (112) using T4 DNA ligase and incubated at 16°C for about 18 hours. The adapter-ligated DNA was then amplified with the primers complementary to the adapter and the transposon inverted repeat sequence using 18 cycles. The PCR product was gel-purified and sequenced at the IU Center for Genomics and Bioinformatics using NextSeq500. Sequencing reads were normalized and subsequently mapped to *B. subtilis* 3610 genome (NCBI accession <u>CP020102</u>) (113) using Bowtie. Between the uninduced and the induced libraries, the distribution of sequencing reads at the TA sites within each gene was compared using Mann Whitney U test. Genes in which reads were statistically underrepresented (*P* value of <0.05) in the induced library compared to the uninduced library were identified. Visual inspection of the transposon insertion profiles was performed with the Sanger Artemis Genome Browser and Annotation tool (https://www.sanger.ac.uk/tool/artemis/) (114).

### P-Galactosidase assays

Three biological replicates of each strain were grown in 3 mL of LB broth at 37°C to an OD_600_ of 0.7–1.0. Two hundred microliters of culture was pelleted and resuspended in 200 μL of Z-buffer (16.1 g Na_2_HPO_4_ • 7H_2_O + 5.5 g NaH_2_PO_4_ • H_2_O + 0.75 g KCl + 0.246 g MgSO_4_ • 7H_2_O in 1 L H_2_O, pH = 7) with 0.27% βME. Four microliters of 10 mg/mL lysozyme was added, and cells were lysed by incubating samples at 30°C for 15 min. In a Greiner-Costar 96-well plate, lysates were diluted to 1:10 in Z-buffer to 200 μL, i.e. 180 μL of Z-buffer and 20 μL of lysates. Three wells had 200 μL of Z-buffer only and served as negative controls. Forty microliters of 4 mg/mL ortho-nitrophenyl-β-D-galactopyranoside (ONPG) in Z-buffer was added to each reaction. The plate was incubated at 30°C for 1 h, and the OD_420_ and OD_550_ of each well were taken every 2 min. The slope of each sample’s ODs over time was derived. The average slope of the three negative control wells was subtracted from the slope of each experimental well. The Miller Units were calculated using the following formula.

### Strain construction

All constructs were either first introduced into the domesticated strain PY79 by natural competence and then transferred to the 3610 background using SPP1-mediated generalized phage transduction (115), or by direct transformation into the competent derivative of 3610, DK104 (116). Detailed descriptions of constructs are in **Supplemental Methods**. All strains used in this study are listed in **Table 1**. All plasmids used in this study are listed in **Supplemental Table S1**. All primers used in this study are listed in **Supplemental Table S2**.

**Table 1:**
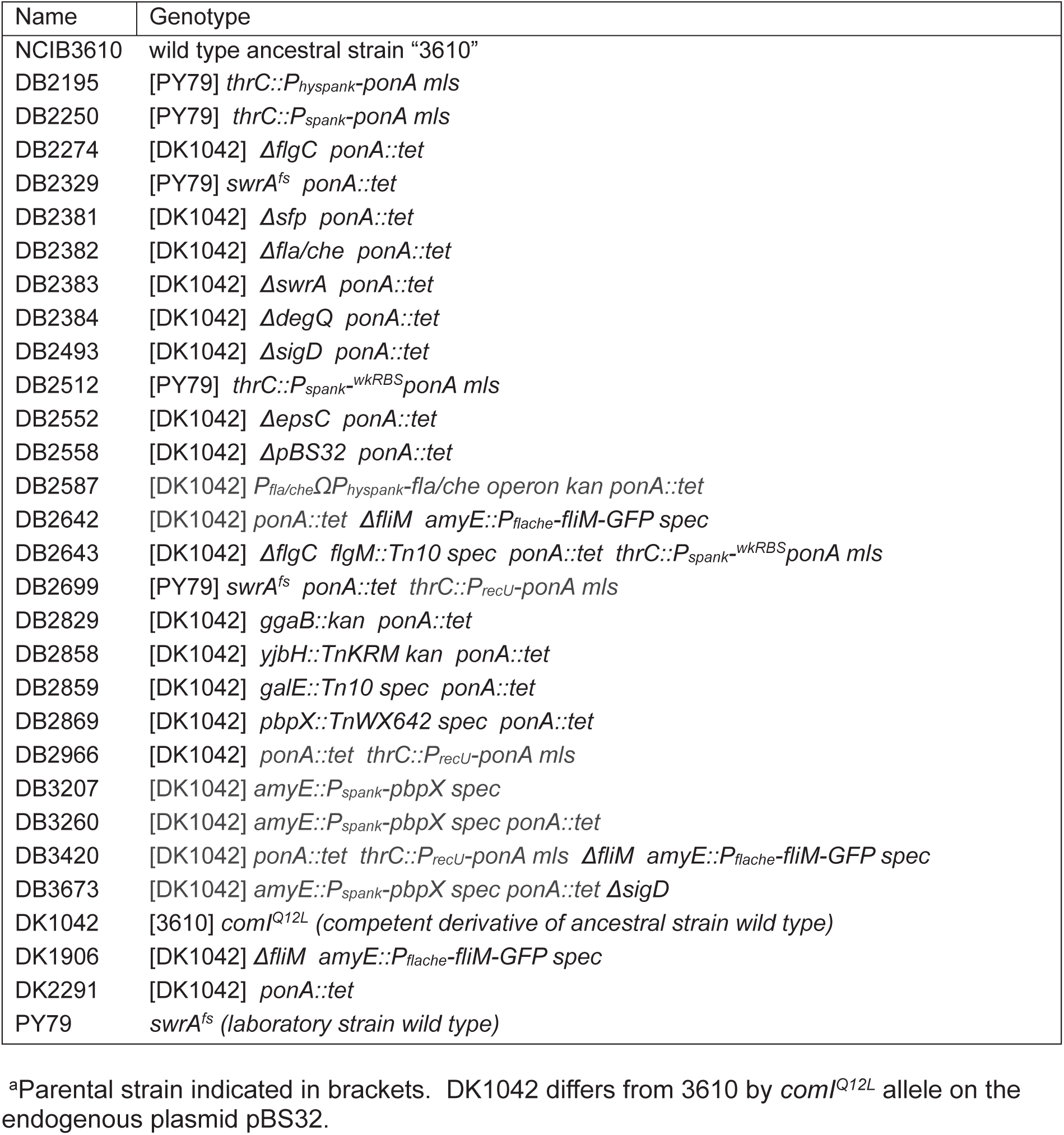
Strains^a^.

## DATA AVAILABILITY

The TnSequencing datasets are available at NCBI Sequence Read Archive (SRA) accession number PRJNA1454591 (https://www.ncbi.nlm.nih.gov/bioproject/1454591)

## ACKNOWLEDGEMENTS

We thank Sid Shaw, Julie Biteen, and Daniel Foust for continued support with flagellar localization and tracking, and Xindan Wang for continued support with transposon sequencing. We thank Rebecca Calvo for construction of the *ponA::tet* allele. We thank Jim Powers and Andras Kun assistance at the Indiana University Light Microscopy Imaging Center (LMIC). Support for this work comes from National Institutes of Health grants R35-GM131783 to DBK and NIH1S10OD024988-01 to the Indiana University Light Microscopy Imaging Center (LMIC).

