## Supplemental Text for "Flagellar toxicity: flagellar synthesis is lytic for *Bacillus subtilis* in the absence of PBP1"

#### SUPPLEMENTAL METHODS

**SPP1 phage transduction.** To 0.2 ml of dense culture grown in TY broth (LB broth supplemented after autoclaving with 10 mM $MgSO_4$ and 100 µM $MnSO_4$), serial dilutions of SPP1 phage stock were added and statically incubated for 15 min at 37˚C. To each mixture, 3 ml TYSA (molten TY supplemented with 0.5% agar) was added, poured atop fresh TY plates, and incubated at 37˚C overnight. Top agar from the plate containing near confluent plaques was harvested by scraping into a 50 ml conical tube, vortexed, and centrifuged at 5,000 x g for 10 min. The supernatant was treated with 25 µg/ml DNase final concentration before being passed through a 0.45 µm syringe filter and stored at 4˚C.

Recipient cells were grown to stationary phase in 2 ml TY broth at 37°C. 0.9 ml cells were mixed with 5 µl of SPP1 donor phage stock. Nine ml of TY broth was added to the mixture and allowed to stand at 37°C for 30 min. The transduction mixture was then centrifuged at 5,000 x g for 10 min, the supernatant was discarded and the pellet was resuspended in the remaining volume. Cell suspension (100 µl) was then plated on TY fortified with 1.5% agar, the appropriate antibiotic, and 10 mM sodium citrate.

***ponA::tet*.** To generate the *ponA::tet* allele, the region upstream of ponA was PCR amplified from 3610 chromosomal DNA with primers 3053/3054. The upstream amplicon was then digested with BamHI/PstI and cloned into the BamHI/PstI sites of pDG1515 carrying a tetracycline resistance cassette flanked by two polylinkers (117). Next, the region downstream of *ponA* was PCR amplified from 3610 chromosomal DNA with primers 3055/3056. The downstream amplicon was digested with EcoRI/XhoI and cloned into EcoRI/XhoI sites of the intermediate plasmid to create pRC18.

***$P_{recU}$-ponA* complementation construct.** The *ponA* complementation construct pDP649 was generated by PCR amplifying the $P_{recU}$ promoter region with primers 8698/8699 and the *ponA* coding sequence with primers 8700/8701, both using 3610 chromosomal DNA as a template. The $P_{recU}$ amplicon was digested with EcoRI/SalI, the *ponA* amplicon was digested with SalI/BamHI and the two products were simultaneously ligated into the EcoRI/BamHI sites of pDG1664 containing a polylinker and an erythromycin (mls) antiobiotic resistance cassette between two arms of the *thrC* gene (118).

**$P_{spank}$-$^{wkRBS}$*ponA* inducible construct.** The inducible *ponA* allele was generated in a series of constructs to find one that was both functional and responsive to IPTG addition. First, the *ponA* gene was PCR amplified using 3610 chromosomal DNA as a template and primers 8576/8577. The amplicon was integrated by isothermal assembly (119) into the HindIII site of pDP150 containing a polylinker downstream of the $P_{hyspank}$ promoter, an *mls* antibiotic resistance cassette, and the *lacI* gene between the arms of the *thrC* gene, to generate plasmid pCD51. The plasmid pCD51 was transformed into PY79 to generate strain DB2195. Next, the $P_{hyspank}$ promoter was weakened by site directed mutagenesis to a $P_{spank}$ promoter by generating two amplicons by PCR using DB2195 chromosomal DNA as a template and primer pairs 6037/9159 and 9160/6032. The two arms were then ligated to each other by isothermal assembly and transformed into PY79 to generate strain DB2250. Finally, the ribosome binding site of *ponA* was weakened with site directed mutagenesis by generating two amplicons using DB2250 chromosomal DNA as a template and primer pairs 6037/8681 and 8680/6032. The two arms were then ligated to each other by isothermal assembly and transformed into PY79 to generate strain DB2512 that served as transduction donor.

**$P_{pbpX}$-*lacZ* reporter.** To generate the $P_{pbpX}$ transcriptional reporter construct pDP667, the region upstream of *pbpX* was PCR amplified from 3610 chromosomal DNA using primers 8943/8944.

The PCR product was digested with EcoRI/BamHI and cloned into the same sites of pDG268 containing a polylinker upstream of the *lacZ* gene, and a chloramphenicol resistance cassette between two arms of the *amyE* gene (120).

***$P_{spank}$-pbpX* inducible construct.** To generate the *pbpX* inducible construct pDP674, the *pbpX* gene was PCR amplified from 3610 chromosomal DNA using primers 8977/8978. The PCR product was digested with SalI/NheI and cloned into the same sites of pDR110 containing a polylinker downstream of the $P_{spank}$ promoter gene, and a spectinomycin resistance cassette between two arms of the *amyE* gene (generous gift of David Rudner, Harvard Medical School).

**Sequence alignment.** The amino acid sequence of PbpX from *B. subtilis* NCIB 3610 was queried against the EggNOG database (Huerta-Cepas et al., 2019) to identify orthologous proteins. Representative ortholog sequences were selected and retrieved in FASTA format. Multiple sequence alignment was subsequently performed using T-Coffee (http://tcoffee.crg.cat/apps/tcoffee/do:regular) The resulting alignment was visualized and color-shaded using the Color Align Sequence tool available through the Sequence Manipulation Suite (https://www.bioinformatics.org/sms2/color_align_cons.html)

**Swarm expansion assay.** Cells were grown overnight at room temperature in LB broth, back-diluted and grown to mid-log phase at 37˚C in LB broth and 1mM IPTG (if applicable), and resuspended to an $OD_{600}$ of 10 in MQ $H_2O$ containing 0.5% India ink. Freshly prepared LB plates containing 0.7% Bacto agar (25 ml/plate) and 1mM IPTG (if applicable) were dried for 10 minutes in a laminar flow hood, centrally inoculated with 10 µl of the cell suspension, dried for another 10 minutes, and incubated at 37˚C for 6 hours. Each strain was done in technical triplicate. The India ink demarcates the origin of the inoculation, and the swarm radius was measured in mm relative to this origin.  For consistency, an axis was drawn on the back of the plate and swarm radii measurements were taken along this axis.

**Table S1: Plasmids**

| plasmid | Genotype |
|---|---|
| pCD51 | *thrC::*$P_{hyspank}$*-ponA mls amp* |
| pDG268 | *amyE::lacZ cat amp* |
| pDG1515 | *tet amp* |
| pDG1664 | *thrC::mls amp* |
| pDP150 | *thrC::*$P_{hyspank}$ *mls amp* |
| pDP649 | *amyE::*$P_{recU}$*-ponA mls amp* |
| pDP667 | *amyE::*$P_{pbpX}$*-lacZ cat amp* |
| pDP674 | *amyE::*$P_{spank}$*-pbpX spec amp* |
| pDR110 | *amyE::*$P_{spank}$ *spec amp* |
| pRC18 | *ponA::tet amp* |

**Table S2: Primers**

| Number | sequence |
|---|---|
| 3053 | AGGAGGGATCCACCATCAAATCGAGCATATGAAGCA |
| 3054 | CTCCTCTGCAGATCTCTTTTCCATTTTTATCATAAATCGT |
| 3055 | AGGAGGAATTCAAACTCATCATCCATTGAAAAAACAAAT |
| 3056 | CTCCTCTCGAGAGCTTCAGCAGGATATTAAATCAATC |
| 6032 | TACATGGTCACCGGAATCAT |
| 6037 | CGCATCCGCTAAATGAACGT |
| 8576 | TTAAGCTTAGTCGACAGCTAGTGATTAACGAAAGGTTGAGATGTT |
| 8577 | AATTAGCTTGCATGCGGCTAGTGTTTAATTTGTTTTTTCAATGGATGATGA |
| 8680 | AAAGGAAAAAAGCGTATGTCAGATCAATTTAACAGCCGT |
| 8681 | ATCTGACATACGCTTTTTTCCTTTTAATCACTAGCTGTCGACTAAGCTTAA |
| 8943 | AGGAGGAATTCGCGGAAGCATTGGTTCGAAGCGG |
| 8944 | CTCCTGGATCCCCATTTTGCACACGTTTTGTTAGAC |
| 8977 | AGGAGGTCGACCGTGTGCAAAATGGAATTTATTTTGG |
| 8978 | CTCCTGCTAGCGTTTGCAGGCCAGTCTTCTTTTC |
| 9159 | CGCTCACAATTCCACACATTATGC |
| 9160 | GCATAATGTGTGGAATTGTGAGC |

**Table S3: Supplemental strains**

| Name | Genotype |
|---|---|
| DB2272 | *ΔflhO ponA::tet* |
| DB2273 | *ΔflhP ponA::tet* |
| DB2291 | *ponA::tet* |
| DB2677 | *ponA::tet thrC::*$P_{recU}$*-ponA mls* |
| DB2698 | *ΔfliT ponA::tet* |
| DB2773 | *ΔmotB ponA::tet* |
| DB2774 | *ΔmotA ponA::tet* |
| DB2775 | *Δhag ponA::tet* |
| DB2776 | *ΔflgL ponA::tet* |
| DB2777 | *ΔflgN ponA::tet* |
| DB2778 | *ΔfliW ponA::tet* |
| DB2779 | *ΔyviE ponA::tet* |
| DB2780 | *ΔlytC ponA::tet* |
| DB2781 | *ΔcwlQ ponA::tet* |
| DB2782 | *ΔlytD ponA::tet* |
| DB2787 | *ΔlytF ponA::tet* |
| DB2788 | *ΔcsrA ponA::tet* |
| DB2795 | *ΔfliD ponA::tet* |
| DB2800 | *ΔyvyF ponA::tet* |
| DB2801 | *ΔyviF ponA::tet* |
| DB2803 | *ΔfliS ponA::tet* |
| DB2804 | *ΔflgK ponA::tet* |
| DB2827 | *ΔyolA ponA::tet* |
| DB2828 | *ΔyolB ponA::tet* |
| DB2829 | *ponA::tet ggaB::kan* |
| DB2859 | *ponA::tet galE::Tn10 spec* |
| DB2869 | *ponA::tet pbpX::TnWX642 spec* |
| DB2870 | *yxkC::kan ponA::tet* |
| DK1042 | *wild type* |
| DS9630 | *ponA::tet* |

#### SUPPLEMENTAL FIGURE LEGENDS

**Supplemental Figure Legend 1. Mutation of *ponA* abolishes swarming motility.** Quantitative swarm expansion measurements on 0.7% LB agar plate over time. Each point is an average of three replicates. The following strains were used to generate this data: WT (DK1042), *ponA* (DS9630), *ponA complement* (DB2677).

**Supplemental Figure Legend 2. The growth defect of a *ponA* mutant is not rescued by mutation of most SigD-activated genes.** Each graph plots the optical density measured over time of cells grown in microtiter dishes with LB broth, shaken at 37°C. Gray lines indicate the wild type and black lines indicate the *ponA* mutant. Pink and blue lines indicate cells doubly mutated for *ponA* and the gene indicated. Pink lines indicate mutations that failed to restore robust growth to the *ponA* mutant while blue lines indicate mutations that restored growth to near wild type levels. All graphs use the ancestral 3610 genetic background. The following strains were used to generate these graphs: WT (DK1042), *ponA* (DB2291), *csra ponA* (DB2788), *cwlQ ponA* (DB2781), *flgK ponA* (DB2804), *flgL ponA* (DB2776), *flgN ponA* (DB2777), *flhO ponA* (DB2272), *flhP ponA* (DB2273), *fliD ponA* (DB2795), *fliS ponA* (DB2803), *fliT ponA* (DB2698), *fliW ponA* (DB2778), *hag ponA* (DB2775), *lytC ponA* (DB2780), *lytD ponA* (DB2782), *lytF ponA* (DB2787), *motA ponA* (DB2774), *motB ponA* (DB2773), *yolA ponA* (DB2827), *yolB ponA* (DB2828), *yviE ponA* (DB2779), *yvyC ponA* (DB2801), *yvyF ponA* (DB2800), *yxkC ponA* (DB2870).

**Supplemental Figure Legend 3. Mutations that disrupt PBPX and the regulatory proteolysis system of Spx improve growth when flagella are induced in the absence of PBP1.** A TnSeq scatter plot depicting the relative fitness advantage of various genes when the fla/che operon is induced in the absence of PBP1. To determine the relative fitness, the transposon insertion count for a particular gene in PBP1 mutant cells induced for the fla/che operon with 1 mM IPTG, divided by the transposon insertion count for the same strain in the absence of inducer, all expressed as the statistical pvalue (Mann-Whitney U test) for that particular ratio. Thus genes to the right of the graph generally improve growth in the absence of PBP1 when the *fla/che* operon is induced and points high on the graph indicate high statitistical confidence. We note however, that the number of insertions in the denominator was low to begin with as certain insertions dominated the pool even when inducer was absent. Thus, strict mathematical assessment of the data can be misleading but we nonetheless include the plots here to aid visualization of the genetic logic underpinning candidate gene selection. A) TnSeq scatter plot highlighting SigD regulated genes as large dots. Blue dots indicate SigD-regulated genes (*flhO* and *flhP*) for which transposon insertion density increased when the fla/che operon was induced whereas yellow dots indicate SigD-regulated genes that did not increase in insertion density. B) TnSeq scatter plot highlighting selected candidates (brown dots) that had a large number of insertions and exhibited an increase in insertion density after induction of the induction of the *fla/che* operon. C) TnSeq scatter plot highlighting genes exhibited an increase in insertion density after induction of the induction of the *fla/che* operon and are involved in the proteolytic restriction of Spx (orange dots). The same data set was used to generate each of the three panels.

**Supplemental Figure Legend 4. The growth defect of a PBP1 mutant can be rescued by mutation of *yjbH*.** Each graph plots the optical density measured over time of cells grown in microtiter dishes with LB broth, shaken at 37°C. Gray lines indicate the wild type and black lines indicate the *ponA* mutant. Pink and blue lines indicate cells doubly mutated for the *ponA* and the gene indicated. Pink lines indicate mutations that failed to restore robust growth to the *ponA* mutant while blue lines indicate mutations that restored growth to near wild type levels. All graphs use the ancestral 3610 genetic background. Each point is the average of three replicates. The following strains were used to generate these graphs: WT (DK1042), *ponA* (DB2291), *ponA galE* (DB2859), *ponA ggaB* (DB2829), and *ponA pbpX* (DB2869).

**Supplemental Figure Legend 5. Protein sequence alignment of PbpX homologs.** Sequences aligned from *Bacillus subtilis* (Bsub_PbpX), *Bacillus amyloliquefaciens* (Bamy_PbpX, WP_088612825.1), *Bacillus cereus* (Bcer_PbpX, WP_193648563.1), *Listeria monocytogenes* (Lmo_EGJ24056, HDI4455274.1), *Staphylococcus aureus* (Sau_FmtA, WP_327785247.1), and *Enterococcus faecalis* (Efae_DR75_2700, WP_010827949.1).

Figure S1

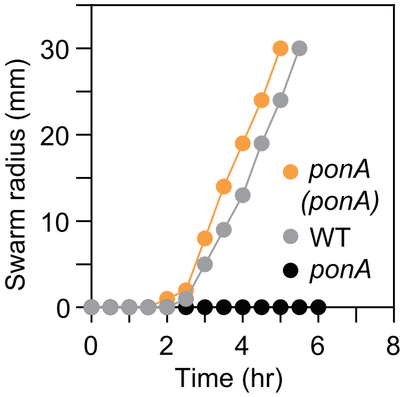

### Figure S2

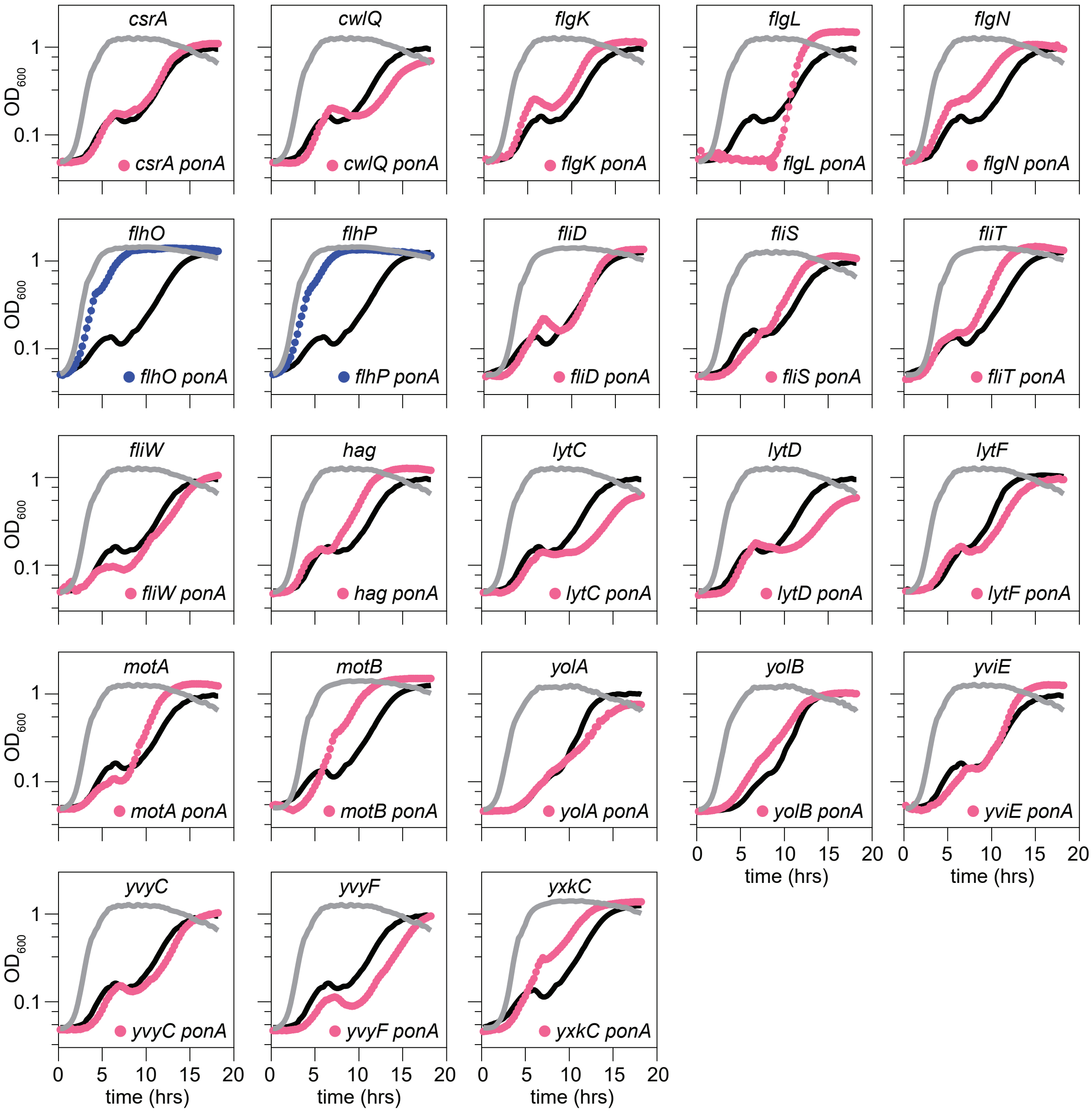
csrA
csrA ponA
cwlQ
cwlQ ponA
flgK
flgK ponA
flgL
flgL ponA
flgN
flgN ponA
flhO
flhO ponA
flhP
flhP ponA
fliD
fliD ponA
fliS
fliS ponA
fliT
fliT ponA
fliW
fliW ponA
hag
hag ponA
lytC
lytC ponA
lytD
lytD ponA
lytF
lytF ponA
motA
motA ponA
motB
motB ponA
yolA
yolA ponA
yolB
yolB ponA
yviE
yviE ponA
yvyC
yvyC ponA
yvyF
yvyF ponA
yxkC
yxkC ponA
OD600
1
0.1
0 5 10 15 20
time (hrs)

Figure S3

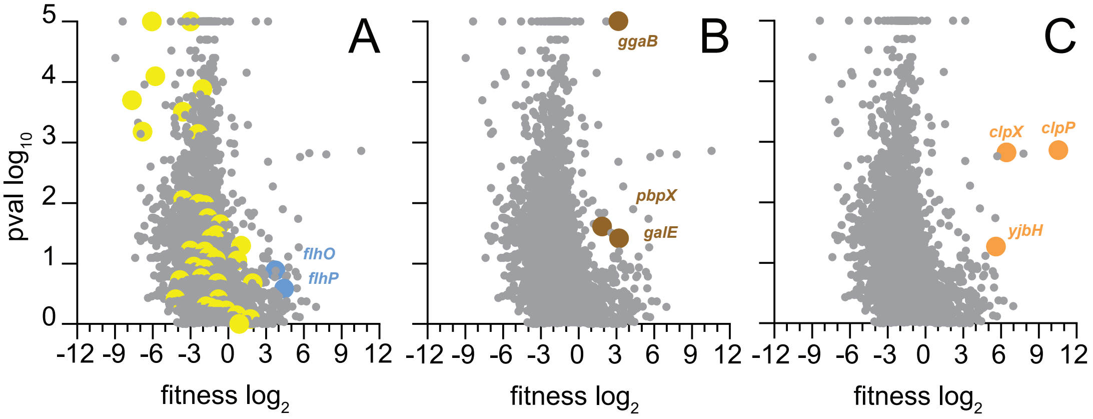
A
B
C
pval log10
fitness log2
fitness log2
fitness log2
flhO
flhP
ggaB
pbpX
galE
clpX
clpP
yjbH
5
4
3
2
1
0
-12 -9 -6 -3 0 3 6 9 12

### Figure S4

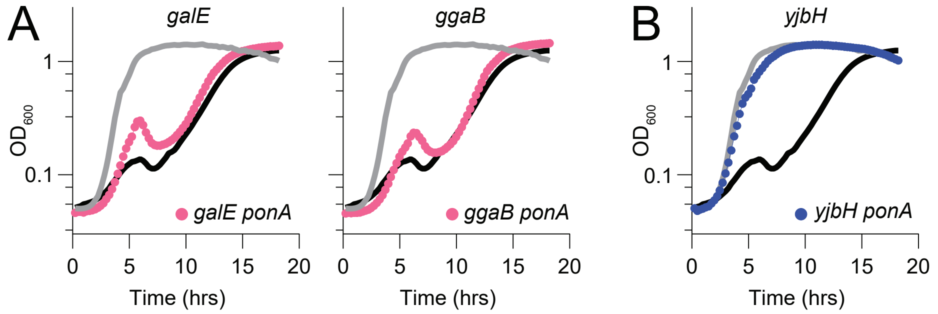

A
B
galE
ggaB
yjbH
1
0.1
OD600
galE ponA
ggaB ponA
yjbH ponA
0
5
10
15
20
Time (hrs)

### Figure S5

```
Bsub_PbpX       MTSPTRRRTAKRRRRKLNKRGKLLFGLLAVMVCITIWN-ALHRNSEENEPSQETAAV-SN-----TD---QKKEV----- 65
Bamy_PbpX       MTSPRRTRINKKRKKKLMKTGGLFLFLLLVTVFIMIWT-VTAHHREKESVKNAPAAQ-SD-----TS---KDQTQ----- 65
Bcer_PbpX       MDQEKNNANGKSRRPIVYIKRTIILVLLFSLVYFMYTK-IVAHNKKEETLKAAAELKKQE-ELKKKEEKKKQEAQKQKEQ 78
Lmo_EGJ24056    MIYEFKSKQ-------------------------------------EQTVSAPTKKK-AATATNPPENAATNEPE----- 37
Sau_FmtA        MKFNK---------VKLVIHACVLLFIIISIAL-------IFHRLQTKTHSIDPIHK-ET-KLSDNEKYLVDRNK----- 57
Efae_DR75_2700  MRKR----HAKKRHGGVN---WLFIVCLLVVIGGSGYLIKTFFFTRDSQVSQESKVV-LE-----ED-----RRS----- 57

Bsub_PbpX       -----KKKT-AKKSEEQIKTV-DRNQKISNYLKEIGFSGTAMIVRNGEIVTNKGFGYADRKHYIQNNPLTSFYVGSSQKA 138
Bamy_PbpX       -----QKKE-KTQKEKTPKSI-DHNKQISNYLQQIGFSGTAMIVKDGKVVLKKGFGYADRKQKIKNNPETSYYIGSSQKA 138
Bcer_PbpX       EEQVKQVQAAEQPAEGPPQEI-NENAQLDQYLQKTGFSGTAVIVKNGKVLLNKGYGMANKEKQVPNNSETTFYIGSISKA 157
Lmo_EGJ24056    -----ETKEPEPVEEKPKETI-VKNQEIDNYLQKIGFSGSALVVRDGKTIIQKGYMYANRDEKVLNTPDTLFYIGSSQKA 111
Sau_FmtA        -----EKVAPSKLKEVYNSKD-PKYKKIDKYLQSSLFNGSVAIYENGKLKMSKGYGYQDFEKGIKNTPNTMFLIGSAQKF 131
Efae_DR75_2700  ----------DNYANLTKEIVAPDSGELDQKIQETNYIGSALIIKDDQVLVNKGYGFANFEKQQANTPNTRFQIGSIQKS 127

Bsub_PbpX       LIATAILQLEEKGKLQTSDPVSTYLPHFPNGQTITLKNLLTHTSGINGHIEGNGAITPDDLIKDIELQGIKRQPG-VWDY 217
Bamy_PbpX       LIATAVLQLEEKGLLQTSDPVSTYLPNFPNGTRITLKNLLNHTSGINGHIEGQGPISPIDLIKDIEYRGIKRQPG-VWDY 217
Bcer_PbpX       FVATAIMQLKDQNKLNVEDTVAKYIPDFPQGNGIQLKHLLTHTSGIPEYEQGTEDISHEELIKRIGKQKRIGSPGEKWKY 237
Lmo_EGJ24056    IIATALLQLEEKGKINTNDPISKYIPDFPNGNKILVKNFMNHTSGIVGRPKLTSNMTPDQIIKAIEKRGIKSQPG-KWDY 190
Sau_FmtA        STGLLLKQLEEEHKININDPVSKYLPWFKTSKPIPLKDLMLHQSGLYKYKSSKDYKNLDQAVKAIQKRGIDPKKYKKHMY 211
Efae_DR75_2700  FTTTLILKAIEEGKLTLDTKLATFYPQIQGAEDITISDMLNMTSGLKLSAMPNNIVTDEEIIQFVKQNTIQVNKG-KYNY 206

Bsub_PbpX       KDSNYSVLAYIIAEVSGEPYEQYIKNHIFKPAGMTHAGFYKT-YEKEPYPAVGYKMEGSK--T---VTPYIPDLSQLYGA 291
Bamy_PbpX       RDSNYSVLAYIVSQVSGESYEQYIKEHVFKPAGMKHAGFYKT-FAKAKYPAVGYKVDESG-SL---VTPHLLDLSQLYGA 292
Bcer_PbpX       SDSNYSILAYIAEKVSGQPLEEYIKQHIFATAGMKHSGFGRE-LEQTRFPSTGYKIVNNN--M---TTPNIPSMSQLYGC 311
Lmo_EGJ24056    LDANYAVVGYLVEKVSGQPLATYLQENIFTPAGMKHTGYYQKDVDGGKNVSTGYKIDENG-VF---QSPKLADLTKLYGA 266
Sau_FmtA        NDGNYLVLAKVIEEVTGKSYAENYYTKIGDPLKLQHTAFYDE-QPFKKYLAKGYAYNSTGLSF---LRPNI--LDQYYGA 285
Efae_DR75_2700  SPVNFVLLAGMLEKMYQRTYQELFNNLYHKTAGLKNFGFYET-LLEQPNNSTSYKWTEDN-SYNQVLSIPAASFAHEFGT 284

Bsub_PbpX       GDIYMSAIDMYKFDQALIDGKLYSQKSYE----KMFTPGSSSTYGMGFYVAPGSYSNHGVMPGFNILNSFSKSGQTIVIL 367
Bamy_PbpX       GDMYMSAEDLYKFDKAIVNGTLLSKESFT----KMFTAGSSAAYGMGFYIDPGSYSNHGVMPGYNILNSFSHSASKYVIL 368
Bcer_PbpX       GDIYTSAHDLYLFNEALFSGKLISKESYN----QMFTAV-KKDYGFGWYVDPGSYSNHGVMPGWNCLNGFSKNGSVYVVL 386
Lmo_EGJ24056    GDISMSTTDMYLFDKALMDKKLISEASLK----KMFTSGSKSTYGMGFYVDSGSYNSHGVVTGWDVSNSVSHTGRTYVIL 342
Sau_FmtA        GNLYMTPTDMGKLITQIQQYKLFSPKITNPLLHEFGTKKYPDEYRYGFYAKPTLNRLNGGFFGQVFT--VYYNDKYVVVL 363
Efae_DR75_2700  GNVDMTTGDLYWYLHQLTSGHLVSTALLQ----KLWTSSQQSSYHGGIYVHDNYLRLHGVEAGQQALVLFSKDMKTGVIL 360

Bsub_PbpX       FSNIQNN--AKLGQVNNKIYQLLNQ--E------ 391
Bamy_PbpX       LSNIQNN--AKLGQVNNRIYQLLNQA-D------ 393
Bcer_PbpX       LSNIQNNI-KSFGKVNNDIYTMLRNIEV------ 413
Lmo_EGJ24056    FSNVQNHI-ASFGKVNNDIYGFLN---K------ 366
Sau_FmtA        ALNVKGNNEVRIKHIYNDILKQNKPYNTKGVIVQ 397
Efae_DR75_2700  LTNCVNP--AKYKELIGSLFHDVTNLTVKF---- 388
```
